# Genomic insights on DNase production in *Streptococcus agalactiae* ST17 and ST19 strains

**DOI:** 10.1101/2020.12.09.418327

**Authors:** Inês Silvestre, Alexandra Nunes, Vítor Borges, Joana Isidro, Catarina Silva, Luís Vieira, João Paulo Gomes, Maria José Borrego

**Affiliations:** Department of Life Sciences, UCIBIO, Nova School of Science and Technology, 2829-516 Caparica, Portugal; National Reference Laboratory for Sexually Transmitted Infections, Department of Infectious Diseases, National Institute of Health, Avenida Padre Cruz, 1649-016 Lisbon, Portugal; Bioinformatics Unit, Department of Infectious Diseases, National Institute of Health, Avenida Padre Cruz, 1649-016 Lisbon, Portugal; CBIOS - Research Center for Biosciences & Health Technologies, Lusófona University of Humanities and Technologies, Campo Grande 376, 1749-024 Lisbon, Portugal; Innovation and Technology Unit, Department of Human Genetics, National Institute of Health, Avenida Padre Cruz, 1649-016 Lisbon, Portugal; Centre for Toxicogenomics and Human Health (ToxOmics), Nova Medical School|Faculdade de Ciências Médicas, Universidade Nova de Lisboa, Campo dos Mártires da Pátria, 1169-056 Lisbon, Portugal

**Keywords:** *Streptococcus agalactiae*, Whole Genome Sequencing, DNases, ST17, ST19

## Abstract

*Streptococcus agalactiae* evasion from the human defense mechanisms has been linked to the production of DNases. These were proposed to contribute to the hypervirulence of *S. agalactiae* ST17/capsular-type III strains, mostly associated with neonatal meningitis. We performed a comparative genomic analysis between ST17 and ST19 human strains with different cell tropism and distinct DNase production phenotypes. All *S. agalactiae* ST17 strains, with the exception of 2211-04, were found to display DNase activity, while the opposite scenario was observed for ST19, where 1203-05 was the only DNase(+) strain. The analysis of the genetic variability of the seven genes putatively encoding secreted DNases in *S. agalactiae* revealed an exclusive amino acid change in the predicted signal peptide of GBS0661 (NucA) of the ST17 DNase(-), and an exclusive amino acid change alteration in GBS0609 of the ST19 DNase(+) strain. Further core-genome analysis identified some specificities (SNVs or indels) differentiating the DNase(-) ST17 2211-04 and the DNase(+) ST19 1203-05 from the remaining strains of each ST. The pan-genomic analysis evidenced an intact phage without homology in *S. agalactiae* and a transposon homologous to TnGBS2.3 in ST17 DNase(-) 2211-04; the transposon was also found in one ST17 DNase(+) strain, yet with a different site of insertion. A group of nine accessory genes were identified among all ST17 DNase(+) strains, including the Eco47II family restriction endonuclease and the C-5 cytosine-specific DNA methylase. None of these loci was found in any DNase(-) strain, which may suggest that these proteins might contribute to the lack of DNase activity. In summary, we provide novel insights on the genetic diversity between DNase(+) and DNase(-) strains, and identified genetic traits, namely specific mutations affecting predicted DNases (NucA and GBS0609) and differences in the accessory genome, that need further investigation as they may justify distinct DNase-related virulence phenotypes in *S. agalactiae*.

## 1. Introduction

*Streptococcus agalactiae* is a major human pathogen which, although present as a commensal bacteria in the gastrointestinal and genitourinary tract, is a leading cause of neonatal morbidity and mortality worldwide, often associated with neonatal meningitis, pneumonia and sepsis (Almeida *et al*., 2017; Remmington and Turner, 2018). The ability of *S. agalactiae* to cause a wide range of disease may be due to an extensive arsenal of virulence factors that contribute to pathogenicity (Remmington and Turner, 2018; Spellerberg, 2000), which includes biofilm formation and the production of DNases (Derré-Bobillot *et al*., 2013; Storisteanu *et al*., 2017; Remmington and Turner, 2018). In fact, DNases would be able to destroy neutrophil extracellular traps (NETs) released by neutrophils upon microbial infection, when they are recruited to the infectious site to restrain bacterial spreading (Brinkmann *et al*., 2004; Brinkmann and Zychlinsky, 2012; Liu *et al*., 2017; Storisteanu *et al*., 2017), as NETs are composed of DNA, histones, proteolytic enzymes and other peptides. However, many pathogens have evolved mechanisms to evade NET-mediated entrapment and killing in what appears to be a widespread strategy to allow pathogen proliferation and dissemination (Liu *et al*., 2017; Storisteanu *et al*., 2017; Remmington and Turner, 2018). DNases can also reduce Toll-like receptor 9 (TLR-9) signaling to dampen the immune response and produce cytotoxic deoxyadenosine to limit phagocytosis (Storisteanu *et al*., 2017; Remmington and Turner, 2018). Streptococcal DNases may be further involved in interacting with other microbial communities through bacterial killing and disruption of competitive biofilms, or be able to control their own biofilm production. In addition, the production of DNases may provide nutrients during colonization and infection by scavenging nucleic acids of neutrophils, as NET formation ultimately leads to cell death. This may explain why many streptococci possess both secreted and cell-wall-anchored DNases. However, the role of DNases during pathogenesis is not completely understood and may be wide ranging, extending beyond the direct interference with the host immune response (Liu *et al*., 2017; Remmington and Turner, 2018).

High nuclease activity by *S. agalactiae* strains was firstly reported in 1980 (Ferrieri *et al*., 1980). Advances in molecular biology, namely the use of whole genome sequencing (WGS), evidenced homology between the DNases of different streptococcal species (Remmington and Turner, 2018). In 2013, Derré-Bobillot and co-authors (Derré-Bobillot *et al*., 2013) identified seven genes putatively encoding secreted DNases in *S. agalactiae* NEM316 (*gbs0153, gbs0382, gbs0609, gbs0661, gbs0712, gbs0885* and *gbs0997*), including the major nuclease (nuclease A) coding gene (*gbs0661*). Their results showed that the loss of NucA, which is secreted and capable of degrading NETs, resulted in reduced *S. agalactiae* NEM316 virulence (Derré-Bobillot *et al*., 2013; Storisteanu *et al*., 2017). Besides, it seems likely that other DNases exist in *S. agalactiae* but have yet to be identified (Remmington and Turner, 2018); as such, recent genome screenings confirm the presence of potential extracellular DNases in mostly all available *S. agalactiae* genome sequences (Storisteanu *et al*., 2017).

*S. agalactiae* capsular-type distribution and predominance could change over time; however, it became clear that strains of certain clonal complexes (CCs) possess a higher potential to cause invasive disease, while other harbor mainly colonizing strains (Shabayek and Spellerberg, 2018). *S. agalactiae* hypervirulent ST17/capsular-type III strains gained special interest due to their strong association with neonatal meningitis and late onset disease of the newborn (Jones *et al*., 2003; Lamy *et al*., 2006; Manning *et al*., 2009; Tazi *et al*., 2010; Bellais *et al*., 2012; Florindo *et al*., 2014; Lu *et al*., 2018; Shabayek and Spellerberg, 2018; Kao *et al*., 2019). These strains display secreted and surface-exposed factors, including DNases (Manning *et al*., 2009; Martins *et al*., 2007; Brochet *et al*., 2006); accordingly, ST17 strains were recently reported as DNase producers while the DNase non-producer phenotype was only found among CC19 strains, which are well adapted to the vaginal mucosa, evidence a poor invasion ability, and were less often related to meningitis (Florindo *et al*., 2018; Shabayek and Spellerberg, 2018).

The aim of the present study was to shed some light on the genetic background of DNase production in *S. agalactiae* strains from different clinical scenarios. For this purpose, we performed a comparative genomic analysis between ST17 and ST19 human strains with different cell tropism (invasive *vs* carriage) and diverse DNase production phenotypes.

## 2. Materials and methods

### 2.1. Strain collection

In this study, a total of 33 clinical *S. agalactiae* strains of human origin (28 colonizing and 5 invasive) were analyzed, including isolates with differential DNase activity determined through prior assays (Florindo *et al*., 2018). The selected strains belong to the collections of the Portuguese National Institute of Health (Instituto Nacional de Saúde Doutor Ricardo Jorge, INSA, IP) and the Institute of Medical Microbiology and Hygiene (IMMH), Ulm University, Germany.

For comparative purposes, *S. agalactiae* reference strains exhibiting distinct DNase activities were used: NEM316 (genotype: III/ ST23) (ATCC 12403, GenBank accession number NC004368); COH1 (genotype: III/ST17) (GenBank accession number AAJR01000000); 2603V/R (genotype: V/ST110) (ATCC BAA-611, GenBank accession number NC004116).

### 2.2. Bacterial culture

Bacteria were grown in Columbia agar supplemented with 5% sheep blood (Biomérieux, Marcy l’Etoile, France) at 5% CO_2_ for 24 h. These cultures were used to proceed to DNase activity assays, antimicrobial susceptibility testing, and to inoculate cultures on fresh Todd Hewitt broth (THB) supplemented with 0.5% yeast extract, which were allowed to incubate without shaking at 37°C with 5% CO_2._ Cell growth was monitored by measuring the optical density at 600 nm (OD_600_). At middle exponential phase (OD_600_=0.2-0.5), 1 ml of bacterial cells was collected by centrifugation (3000 rpm, 10 min), resuspended in 200µl of PBS and immediately stored at −20°C, before further DNA extraction.

### 2.3. DNase activity, qualitative assays

DNase production phenotype was assessed qualitatively by growing *S. agalactiae* strains on DNA-methyl green agar plates (Oxoid, Basingstoke, England), that allowed the identification of DNase production. The results were interpreted after 24 h of incubation at 37°C in an atmosphere of 5% CO_2_. Strains were considered DNase producers (DNase(+)) when displaying transparent halos around colonies of *S. agalactiae* in DNA-methyl green agar plates. *S. agalactiae* strains COH1 and NEM316 were used as positive controls and 2603V/R as a negative control (DNase(-)).

### 2.4. DNase activity, semi-quantitative and quantitative assays

In order to confirm the results of the qualitative tests, which pose challenges for interpretation based on visualization, a semi-quantitative assessment of the activity of the DNases was further performed according to the described by Sumby and co-authors (Sumby *et al*., 2005), with modifications. Briefly, *S. agalactiae* strains were inoculated in 5 ml of THB supplemented with 0.5% yeast extract, culture supernatant from stationary phase was centrifuged (10 min., 3000 rpm), syringe filtrated (0.2 µm), before incubation at 37°C with 1 µg of double stranded DNA (*atr* amplicon, Jones *et al*., 2003) in the presence of 1x M buffer (Roche). Filtrated supernatants (10 µl) were incubated for 1h, 2h, 4h and “overnight” (∼ 17h), in order to evaluate the digestion of DNA in a final volume of 50 µl. Nuclease reaction was stopped by adding EDTA (0.5 M, pH 8.0) at 4°C. DNA digestion was analyzed by electrophoresis in 1% agarose gel. A negative control consisting on a reaction mixture without supernatant was used in all experiments.

In order to confirm the results of production / non-production obtained by the methods previously described, a quantitative evaluation of the activity of DNases was further performed for a selection of *S. agalactiae* strains. The quantitative DNase assays were performed as described above and the amount of dsDNA present in each sample was determined by measuring at the same time points of incubation, using Quant-iT^™^ PicoGreen^™^ dsDNA assay kit (Thermo Fisher Scientific, USA), according to manufacturer’s instructions. Briefly, 1 ml of 1x Quant-iT PicoGreen was added to an equal volume of each supernatant, previously diluted in 1x TE buffer. After 5 minutes of incubation at room temperature, in the dark, we proceeded to the fluorescence measurement (Fluorimeter – Anthos Zenith 3100) in 96 well microtiter plates (Corning 96 Well Clear Flat Bottom Polystyrene Black TC, Costar, USA). To calculate the concentration of the remnant DNA in each sample, a standard curve with four solutions of phage Lambda DNA of known concentrations (1, 10, 100 and 1000 ng/ml) was used. Each obtained fluorescence value was subtracted of the blank solution (PicoGreen 1x + 1x TE buffer at a ratio of 1:1) fluorescence value. *S. agalactiae* strains COH1 and NEM316 were used as positive controls and 2603V/R as a negative control.

### 2.5. Nucleic acid isolation

Genomic DNA was extracted as previously described (Florindo *et al*., 2012), with minor changes. Briefly, bacterial cells were subjected to a high-speed centrifugation (14000 rpm) for 10 min at 4°C. The pellet was digested for 2 h at 37°C with 200 µl of Tris-EDTA buffer, pH 8.0, containing 10 U mutanolysin (Sigma-Aldrich, St. Louis, USA) and 15 mg/ml lysozyme (Sigma-Aldrich, St. Louis, USA) before treatment with 10 mg/ml proteinase K (Roche, Penzberg, Germany). Subsequently, DNA was extracted using the NucliSENS® EasyMag® (BioMérieux, Marcy l’Etoile, France) or the Isolate II Genomic DNA kit (Bioline, Tennessee, USA), according to the manufacturer’s instructions. DNA quality, purity and concentration were evaluated by agarose gel electrophoresis, spectrophotometry (NanoDrop, ThermoFisher Scientific, Massachusetts, USA) and fluorimetry (QubitTM, ThermoFisher Scientific, Massachusetts, USA), respectively. Then, DNA samples were sent to sequencing in Innovation and Technology Unit, at the Portuguese National Institute of Health.

### 2.6. Whole genome sequencing, de novo assembly and annotation

Previously selected *S. agalactiae* ST17 and ST19 strains (n=33) were all subjected to WGS, as previously described (Pelerito *et al*., 2020). Briefly, high-quality DNA samples were subjected to dual-indexed Nextera XT Illumina library preparation (Illumina, USA), prior to cluster generation and paired-end short-read high throughput sequencing (2×150bp or 2×250bp) on an Illumina MiSeq or NextSeq550 equipment (Illumina, USA), according to the manufacturer’s instructions. Genomes were *de novo* assembled using the INNUca v3.1 pipeline (https://github.com/B-UMMI/INNUca) (Llarena *et al*., 2018), an integrative bioinformatics pipeline that consists of several integrated modules for reads QA/QC, *de novo* assembly and post-assembly optimization steps. Briefly, after reads’ quality analysis using FastQC v0.11.5 (http://www.bioinformatics.babraham.ac.uk/projects/fastqc/) and cleaning with Trimmomatic v0.36 (http://www.usadellab.org/cms/?page=trimmomatic) (Bolger *et al*., 2014), genomes were *de novo* assembled with SPAdes 3.10 (http://bioinf.spbau.ru/spades) (Bankevich *et al*., 2012), and subsequently improved using Pilon v1.18 (Walker *et al*., 2014).

Some samples were also subjected to long-read nanopore sequencing (Oxford Nanopore Technologies-ONT) in order to increase the contiguity of genome sequences. Briefly, genomic libraries were prepared using the Rapid Sequencing kit (ONT), according to the manufacturer’s protocol. A MinION device (ONT) USB-connected to a laptop computer with i7 Intel processor and 16 GB RAM, was used to sequence the libraries in two R9.4 type flow cells. The MinKNOW software (v19.06.8; ONT) was used to configure run parameters, data acquisition and real-time base-calling (Guppy v3.0.7; ONT). The quality of the MinION reads was assessed using MinIONQC (https://github.com/roblanf/minion_qc). Canu 1.9 was used for error correction of reads, trimming (removing adapters and breaking chimeras) and assembly with genome size of 2.1 Mbp as input (Koren *et al*., 2017). Curation of Canu assemblies was performed using a parallel mapping strategy: 1) of MinION trimmed reads with Minimap2 (https://github.com/lh3/minimap2) (Li, 2018), and 2) of Illumina trimmed reads with Snippy v4.6 (https://github.com/tseemann/snippy) software, followed by polishing with Pilon v1.18. The later was ran for multiple iterations until the polished genomes remained unchanged.

Draft genome sequences were annotated with both RAST server v2.0 (http://rast.nmpdr.org/) (Aziz *et al*., 2008) and Prokka v1.12 (Seemann, 2014). For simplification purposes, NEM316 locus tag (GenBank accession number NC004368.1) was adopted to designate all gene hits identified in the subsequent analyses.

### 2.7. Core-genome single nucleotide variant (SNV)-based analysis

The genetic relatedness among strains was evaluated by a reference-based mapping strategy using Snippy v4.6 software. For each isolate, quality improved Illumina reads were mapped against a representative draft short-read-only assembled genome (2211-04 was used as mapping reference for all 33 strains as well as for ST17 cluster, while 1203-05 was used as mapping reference for ST19 cluster). SNV calling was performed on variant sites that filled the following criteria: *i)* minimum mapping quality of 20; *ii)* minimum number of reads covering the variant position ≥10; and *iii)* minimum proportion of reads differing from the reference of 90%. Core-SNVs were extracted using Snippy’s core module (*snippy-core*). All putative SNVs/indels were carefully inspected and confirmed using the Integrative Genomics Viewer (IGV) v2.8.2 (http://software.broadinstitute.org/software/igv/) (Robinson *et al*., 2011). MEGA7 software (http://www.megasoftware.net) (Kumar *et al*., 2016) was applied to calculate matrices of nucleotide distances and perform phylogenetic reconstructions over the obtained core-genome SNV alignment by using the Neighbor-Joining method (Saitou and Nei, 1987) with the Maximum Composite Likelihood model to compute genetic distances (Tamura *et al*., 2004) and bootstrapping (1000 replicates) (Felsenstein, 1985).

### 2.8 Extended genetic diversity analysis

The PubMLST online platform (http://pubmlst.org/) was used for *in silico* Multilocus Sequence Typing (MLST). For strains for which the *de novo* assembled genome could not be obtained (n=9) (since did not pass FastQC module QA/QC), MLST locus sequences were retrieved by a read-based mapping approach. Alleles and sequence types (STs) not previously described were deposited in the *S. agalactiae* MLST database. CCs were defined using eBurst analysis (http://eburst.mlst.net/v3/mlst_datasets/) (Feil *et al*., 2004).

The occurrence and structure/diversity of CRISPR-Cas systems (clustered regularly interspaced short palindromic repeats/CRISPR-associated proteins) was predicted for all assembled genomes (n=12 for ST17 and n=12 for ST-19) with both CRISPRCasFinder (http://crisprcas.i2bc.paris-saclay.fr/) and CRISPRone (http://omics.informatics.indiana.edu/CRISPRone/) (Zhang and Ye, 2017) web tools (both accessed in May 2020). Only intact elements with the highest confidence score level were considered as legitimate hits. The occurrence and structure of each hit CRISPR-Cas system was confirmed by RAST annotation and reads-based mapping.

In order to obtain an overview on the repertoire of putative virulence genes carried by each strain, a vast database was constructed, enrolling: *i)* previously identified virulence determinants (Glaser *et al*., 2002, Da Cunha *et al*., 2015) and *ii)* genes identified after querying draft genome sequences against the Virulence Factors Database (VFDB; http://www.mgc.ac.cn/VFs/; last updated at 30^th^ September 2020) (Chen *et al*., 2005), using the VFanalyzer platform (accessed in October 2020). The presence/absence and/or variability of each putative identified hit was confirmed by an assembly-free strategy using Snippy 4.6, while BLASTp against the non-redundant (nr) protein sequences database was used to assess the potential protein function. The identified hits were then submitted at to PubMLST online platform for allele determination.

As a means to potentiate isolate discrimination, assemblies were also analyzed using PHASTER (https://phaster.ca/) (Arndt *et al*., 2016) to determine the presence of prophages. Only hits with intact phages were considered for further analysis, given that the detection of phage fragments could result from the assembly process hampering the distinction between presence of intact phages and phage remnants that were excised during the isolate evolutionary process (for which the biological importance is even more uncertain). The identification of other mobile genetic elements (MGEs) was carried out by manual inspection of annotated sequences, and further confirmed through BLASTn and BLASTp searches against the nr and WGS databases. In particular, the presence of plasmids was assessed through PlasmidFinder 2.0 (http://cge.cbs.dtu.dk/services/PlasmidFinder/), using default parameters with both trimmed reads and assemblies.

Finally, global pan-genome analyses were carried out for all assemblies using Roary v3.8.0 software (Page *et al*., 2015) with a BLASTp minimum percentage of identity set to 95% and without splitting paralogs. In order to obtain referential locus tags, gene hits found to be present/absent exclusively in ‘exception strains’ for DNase activity, were confirmed through reads’ mapping and Blastn search (90% query coverage with 70% identity) against publicly annotated *S. agalactiae* closed or draft genomes.

### 2.9. DNases’ structural characterization

For all DNases, the respective predicted protein sequences were searched for the presence of domains and/or motifs indicative of nucleases. SignalP-5.0 Server (https://www.cbs.dtu.dk/services/SignalP/) was used to check the presence of signal peptides. Putative structures were predicted using SWISS-MODEL (http://swissmodel.expasy.org/).

### 2.10. Data availability

All raw sequence reads used in the present study were deposited in the European Nucleotide Archive (ENA) under the study accession number PRJEB41294 (Table S1).

## 3. Results

### 3.1. DNase activity assays

Of the total 33 clinical *S. agalactiae* selected strains, 15 belonged to ST17 and 18 belonged to ST19. Based on qualitative and semi-quantitative assays, all *S. agalactiae* ST17 strains, with the exception of the clinical strain 2211-04, were found to display DNase activity. The opposite scenario was observed for ST19, where DNase activity was only seen for one (1203-05) out of the eighteen ST19 clinical strains. The presence or absence of DNase activity was further assessed by quantitative assays for a selection of *S. agalactiae* strains from both STs (Figure 1). Confirming the previous observations, the ST17 2211-04 clinical strain was the only ST17 strain lacking DNase activity (congruent with the negative control 2603V/R), while ST19 clinical strains (with exception of 1203-05) showed no ability to degrade DNA, whose amount remained nearly constant after 17h. Interestingly, two distinct DNA digestion profiles were observed for the remaining ST17 clinical strains, when compared with the two positive controls, *S. agalactiae* COH1 and NEM316. While some of them mirrored the medium DNA digestion of COH1, others exhibited an almost total DNA digestion (<10% DNA remained after 17h), reflecting a high production of extracellular DNases like reference NEM316. On the other hand, the ST19 1203-05 clinical strain appeared isolated, revealing only a weak DNase activity.

**Figure 1.**
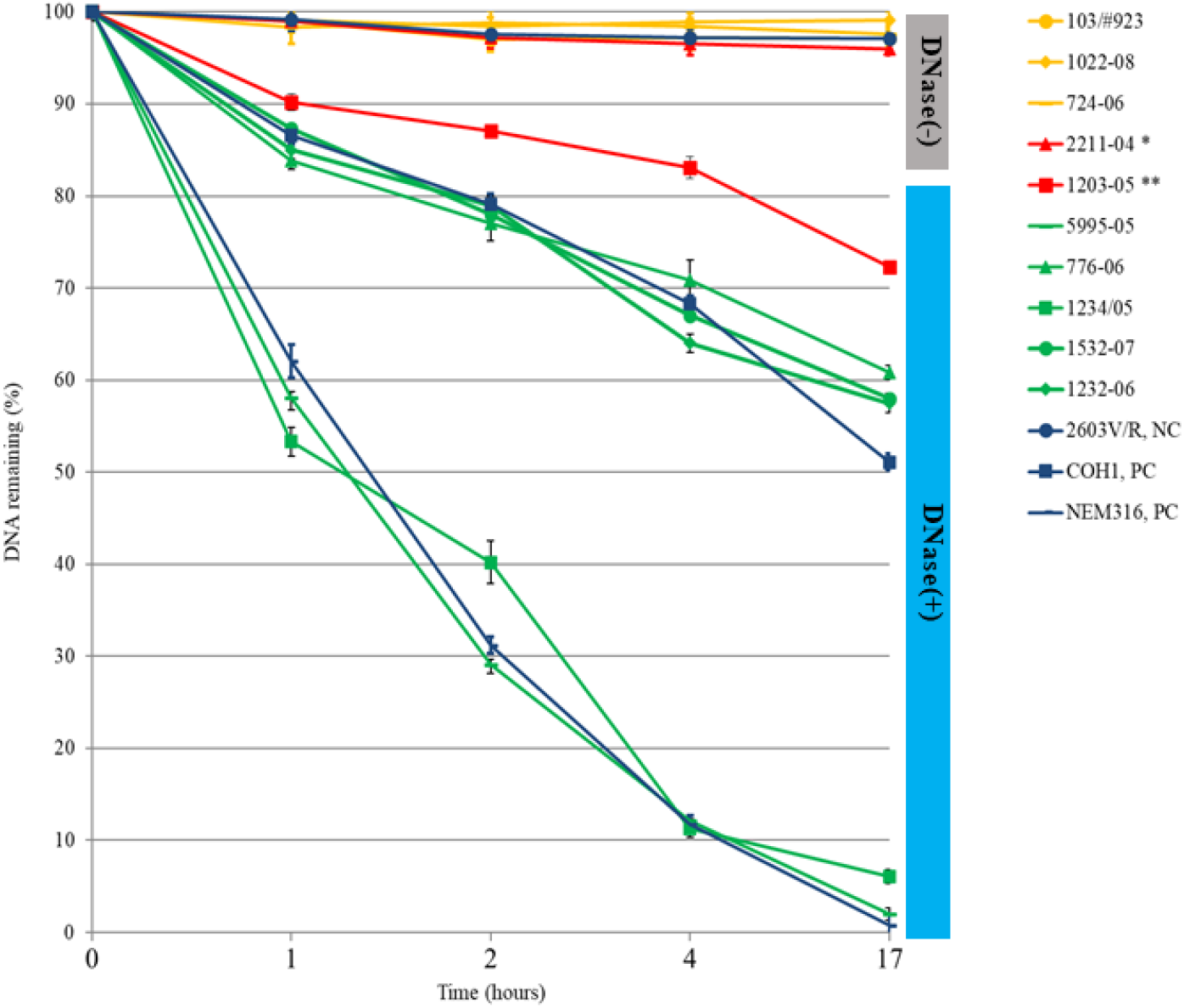
Quantitative DNase assays displaying differential DNase activity between *S. agalactiae* strains over time. *S. agalactiae* culture supernatant (10 μl) incubated with 1 μg of DNA (amplicon *atr*) for 1h, 2h, 4h, overnight at 37°C. Fluorescent PicoGreen dye (Invitrogen) was used to quantify the dsDNA. The graphic displays the mean values of the experiment performed in duplicate. Error bars show ± standard deviation. DNase(-) ST19 strains are shown in yellow, while DNase(+) ST17 strains are depicted in green. The two exceptions are displayed in red, with * representing the DNase(-) ST17 strain and ** representing the DNase(+) ST19 producer. All controls are shown in dark blue (PC, Positive Control; NC, Negative Control).

### 3.2. Strains’ phylogenetic relationship

Considering the observed DNase activity exceptions within each ST, we first assessed the genetic relationship between 2211-04 and 1203-05 and the remaining clinical strains. We observed that the core-genome SNV-based phylogenetic analysis clearly segregates strains based on each ST (Figure 2), resulting in a strong two major-branches tree, where neither ST17 2211-04 nor ST19 1203-05 strains showed up in isolated branches. Indeed, clinical strains from ST17 displayed a mean of 163.2±5.3 core-SNVs among them but had a mean distance of 7342.2±33.2 core-SNVs to ST19 strains, which differ themselves by a mean of 466.5±7.3 core-SNVs (Figure 2A). Strains’ relatedness within each individual ST was also explored by maximizing the core-genome under evaluation (Figure S1). Despite only a slight increase in the number of core-SNVs was observed (mean core-SNVs of 258.9±4.8 within ST17 and mean core-SNVs of 526.2±5.2 within ST19), we were able to identify the clinical strains genetically closest to ST17 2211-04 and ST19 1203-05 strains. For ST17, 2211-04 form an independent phylogenetic clade with 1232-06 (differing by 83.0±9.0 core-SNVs), while for ST19, 1203-05 was found to be genetically closest related to 461-05, 724-06, 1434-05, 1237-07 and 225-06 strains (differing only by a mean of 41.8±5.7 core-SNVs from these strains).

**Figure 2.**
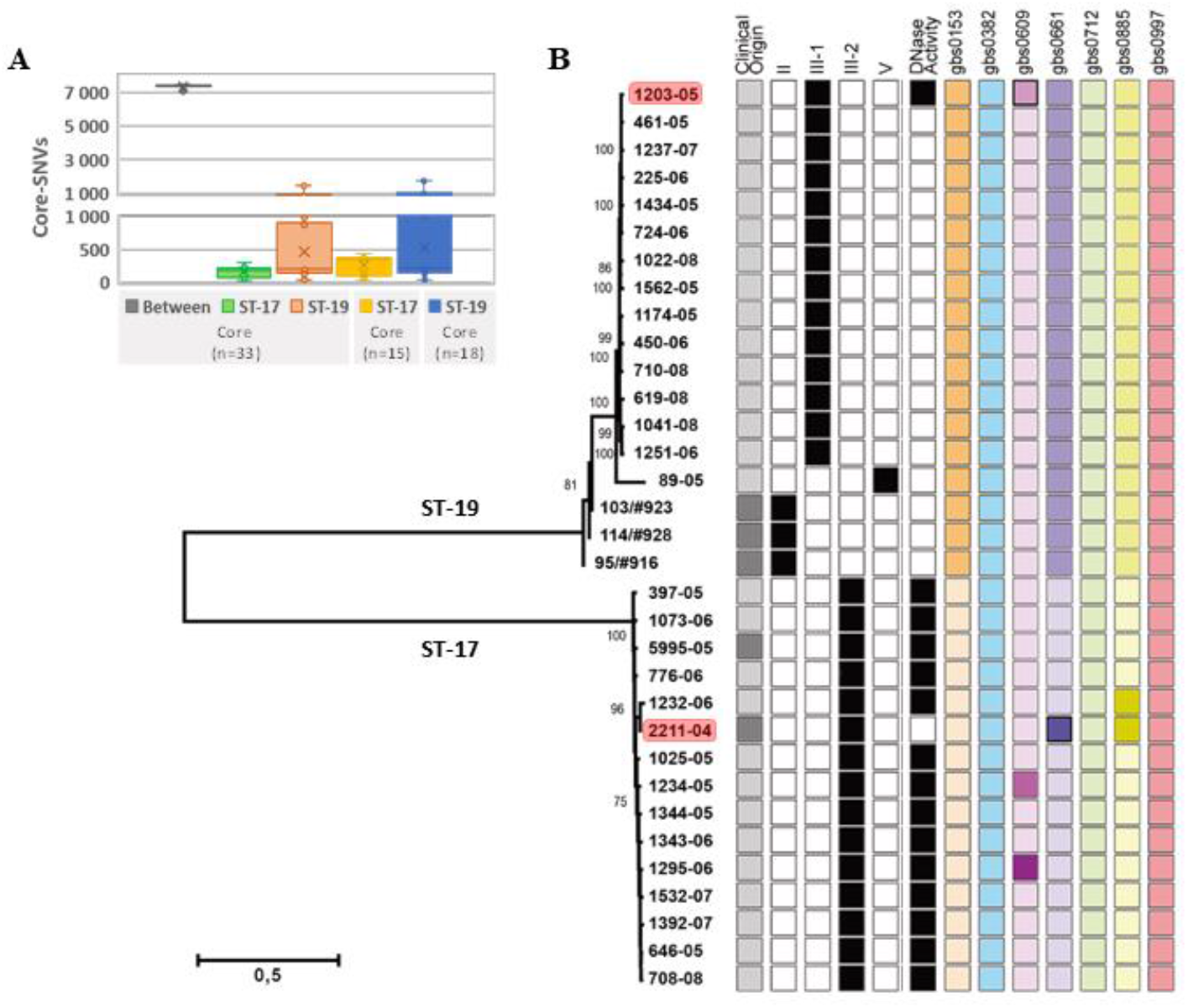
DNase-based genotypes and phenotypes. **A**. Box Plots depicting the number of SNVs observed within the overall population and within each ST. The three left plots are based on a core-genome enrolling all the 33 strains, whereas the two right plots are relative to a core-genome enrolling solely same-ST strains (n=15 for ST17 and n=18 for ST19), in order to maximize the genome shared for each sub-set of strains. The horizontal line within each box marks the median, while the cross (“x”) represents the medium value. **B**. The core-genome SNP-based phylogenetic tree was reconstructed using the Neighbor-Joining method (Saitou and Nei, 1987) with the Maximum Composite Likelihood model (Tamura *et al*., 2004). Bootstraps values are shown next to the respective nodes. Colonization strains are represented by grey light squares, while strains from invasive infection are in grey dark squares. For each isolate, the black squares represent the respective capsular-type as well as the presence of DNase activity (illustrated in Figure 1). Both the DNase(-) ST17 and the DNase(+) ST19 strains are highlighted in red. For each DNase gene, the different alleles found for each clinical strain are represented by distinct color tones (proximal color tones do not reflect genetic proximity). Genes were designated according to NEM316 genome (GenBank accession number NC004368.1).

### 3.3. Genetic variability of DNase genes

We then proceeded with a more focused analysis where the potential genetic basis behind the observed DNase activity was evaluated. All known loci putatively encoding secreted DNases in *S. agalactiae* were analyzed. While some genes were found to be conserved among all strains (the paralogs *gbs0382, gbs0712* and *gbs0997*), or to fully separate strains according to ST (*gbs0153*), specific alleles were exhibited by ST19 1203-05 and ST17 2211-04 clinical strains in *gbs0609* and *gbs0661*, respectively (Figure 2B), which may contribute to their exception character regarding DNase activity. For instance, the weak DNase(+) ST19 1203-05 strain was found to harbor an exclusive missense alteration in *gbs0609*, corresponding to the substitution of a negative-charged amino acid by a positive-charged one (Glu261Lys). On the other hand, for nuclease A coding gene (*gbs0661*), an exclusive non-synonymous alteration (Thr10Ala) affecting the predicted signal peptide was exhibited by the DNase non-producer ST17 2211-04 strain. When compared to the well-studied *gbs0661* of DNase(+) NEM316, all 33 *S. agalactiae* strains additionally displayed one non-synonymous (T296C; Pro99Leu) and two silent (G99A and G228A) substitutions, despite all DNase(-) ST19 strains harbored an extra silent alteration (A423G). Curiously, the replacement of the proline residue by a leucine fall within β-strand 4 of NucA, according to its predicted structure for NEM316 (Moon *et al*., 2014), which together with β-strands 9 and 10 constitute the deepest recess of the nuclease active site (Figure S2). Despite the detected amino acid replacements may hypothetically explain the differences in DNase activity observed in the present study, to our knowledge they have not yet been subjected to any mutagenesis assays, so their role in the enzyme secretion/activity remains unknown.

### 3.4. Core-genome analysis

We then searched for other genetic aspects that could explain the observed DNase activity exceptions. Globally, a total of 35 core-SNVs/indels were found to differentiate DNase(-) ST17 2211-04 from all DNase(+) ST17 strains (Table 1). With exception of five core-SNVs falling within non-coding sequences, variants (17 non-synonymous, 11 synonymous, one frameshift deletion and one conservative in-frame insertion) were found to affect genes mostly related to metabolism (n=13) as well as to cellular processes and signaling (n=11), such as those encoding the ATP-dependent Clp protease ATP-binding subunits ClpC and ClpX, the molecular chaperone DnaK, and the class I S-adenosylmethionine (SAM)–dependent methyltransferase. Regarding the core-SNVs affecting non-coding regions, none of them was found to interrupt any annotated regulatory element (like ribosome binding sites and transcriptional terminators, for instance). Nevertheless, one SNV apparently falls within a region similar to a transposase C-terminal end (pseudogene).

**Table 1.**
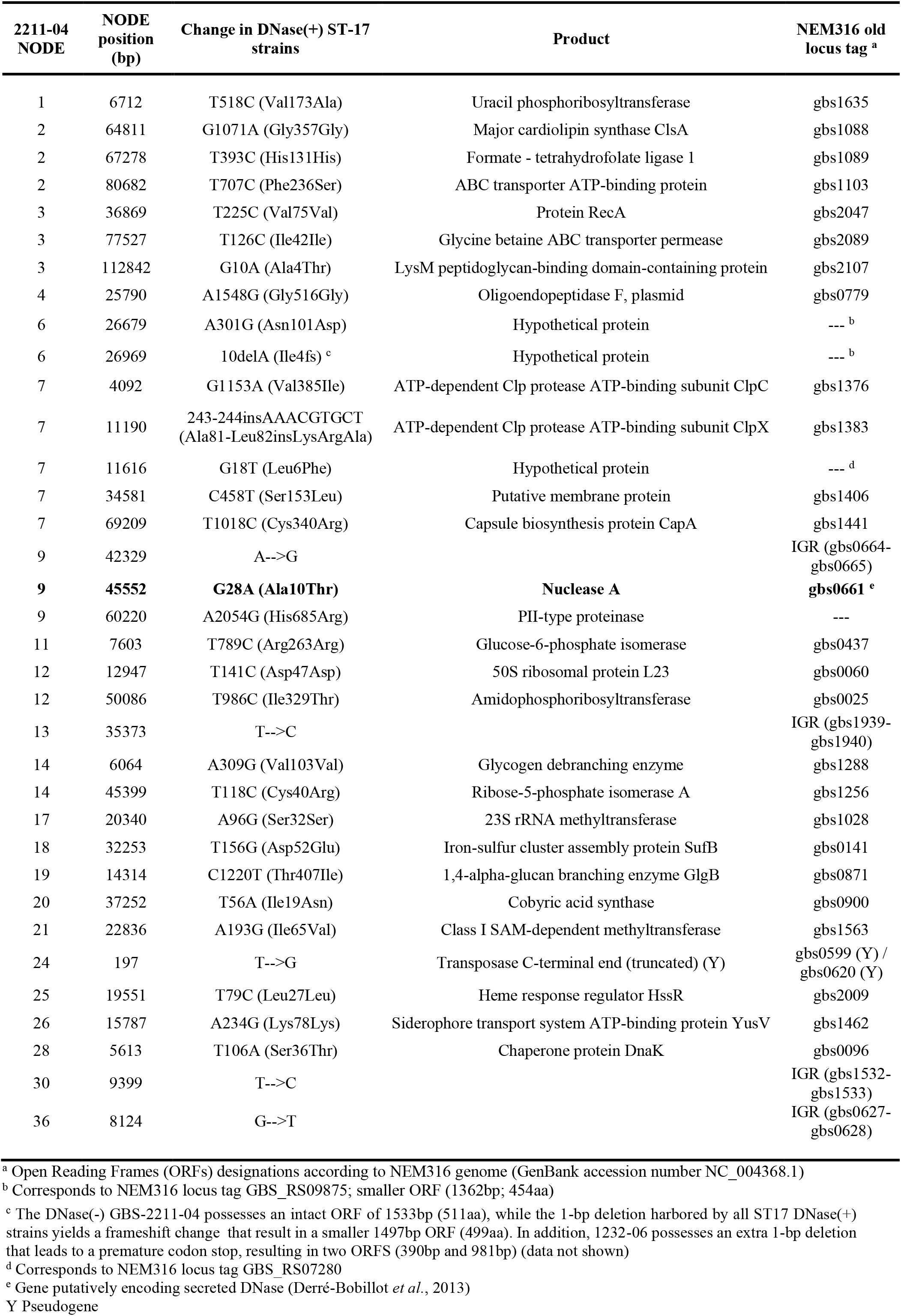
Core-SNVs/indels between DNase(-) 2211-04 and all DNase(+) ST17 strains.

On the other hand, for ST19, 24 core-SNVs/indels were found to discriminate the weak DNase(+) 1203-05 from all DNase(-) ST19 strains (Table 2). Of the 20 core-SNVs occurring in coding sequences, approximately two thirds originate missense variants, and affect genes encoding hypothetical proteins as well as genes related to bacterial stress response, defense mechanisms or metabolism. From these, we highlight two genes encoding the GTPase Obg and the excinuclease ABC subunit B that are related to bacterial stress response and DNA repair (Bonventre *et al*., 2016; Grossman and Yeung, 1990). Interestingly, one core-SNV (G297A) involves the exclusive gain for 1203-05 of a premature stop codon (at 99aa) in the TetR/AcrR family transcriptional regulator gene, likely producing two shorter proteins (99aa and 56aa instead of the original 183aa). Two insertions (of 1bp and 18bp) occurring in genes coding for a nucleoid-associated protein and an hypothetical protein were also found to differentiate all ST19 DNase(-) strains from the weakly DNase(+) 1203-05, yielding proteins with distinct length sizes. Regarding the core-SNVs falling in non-coding sequences, one occurred in the ribosome binding site (RBS) of a *gbs1083* homolog. While all ST19 DNase(-) possess a GGCAGG RBS sequence, the weakly DNase(+) displays an exclusive GGCAGA sequence, which may hypothetically influence the ribosome binding and, consequently, the protein (threonine/serine exporter family protein) expression.

**Table 2.** Core-SNVs/indels between DNase(+) 1203-05 and all DNase(-) ST-19 strains.

| 1203-05<br>NODE | NODE<br>position<br>(bp) | Change in DNase(-) ST-19 strains | Product | NEM316 old<br>locus tag <sup>a</sup> |
| --- | --- | --- | --- | --- |
| 2 | 5182 | G250A (Glu84Lys) | Hypothetical protein | gbs0488 |
| 2 | 14321 | T663G (Asn221Lys) | Hypothetical protein | gbs0496 |
| 3 | 42682 | 264-265insA (Leu89fs) <sup>b</sup> | Nucleoid-associated protein | gbs1791 <sup>b</sup> |
| 4 | 37354 | T926C (Leu309Ser) | Streptococcal histidine triad protein<br>precursor (C-terminal part) | gbs1924 (Y) |
| 4 | 57207 | A591G (Ala197Ala) | NAD-dependent malic enzyme | gbs1906 |
| 5 | 75372 | A923T (His308Leu) | Zinc protease | gbs2113 |
| 6 | 85300 | T-->C |  | IGR (gbs1083-gbs1084) |
| 6 | 102417 | C912T (Pro304Pro) | Phosphoglucomutase | gbs1100 |
| 8 | 62062 | T-->C |  | IGR (gbs0923-gbs0924) |
| 9 | 52087 | T317C (Leu106Pro) | YlbF/YmcA family<br>competence/transcriptional regulator | gbs1446 |
| 10 | 5774 | T434C (Phe145Ser) | FMN-dependent NADH-azoreductase 1 | gbs0259 |
| 10 | 47782 | G1511A (Arg504His) | Ribonuclease Y | gbs0295 |
| 10 | 78357 | A33C (Ile11Ile) | Ribosome hibernation promotion factor | gbs0326 |
| 15 | 42542 | T305C (Ile102Thr) | GTPase Obg | gbs1537 |
| 15 | 49967 | C1555G (Leu519Val) | UvrABC system protein B | gbs1531 |
| 16 | 16085 | A220G (Ser74Gly) | putative FMN/FAD exporter YeeO | gbs0056 |
| 16 | 30300 | T795C (Pro265Pro) | N5-carboxyaminoimidazole ribonucleotide<br>synthase | gbs0043 |
| 19 | 147 | T-->C |  | IGR (gbs1395-gbs1396) |
| 19 | 13854 | A297G (Ter99Trpext*) | TetR family transcriptional regulator | gbs1381 |
| 20 | 38268 | 2892-2909dupAGCACGTAAAGCTGAAGA<br>(Glu970-Gly971insAlaArgLysAlaGluGlu) <sup>c</sup> | Hypothetical protein | --- |
| 20 | 50146 | A630C (Lys210Asn) | Protein DedA | gbs0662 |
| 22 | 7144 | A324G (Ala108Ala) | Cystathionine beta-lyase | gbs1283 |
| **30** | **10716** | **A781G (Lys261Glu)** | **Endonuclease** | **gbs0609 <sup>d</sup>** |
| 45 | 549 | T-->C |  | IGR (gbs0874-gbs0875) |

<sup>a</sup> Open Reading Frames (ORFs) designations according to NEM316 genome (GenBank accession number NC\_004368.1)
<sup>b</sup> The 1bp insertion leads to a premature stop, yielding a smaller ORF in all ST19 DNase(-) strains and NEM316
<sup>c</sup> The 18bp in-frame insertion yields a longer ORFs in all ST19 DNase(-) strains
<sup>d</sup> Gene putatively encoding secreted DNase (Derré-Bobillot *et al.*, 2013)
Y Pseudogene

### 3.5. Pan-genomic analyses

We also aimed to identify accessory genes with the potential to discriminate the two strains exhibiting DNase activity exceptions with the respective STs. As such, we performed a pairwise comparison of gene content between DNase(-) ST17 2211-04 and the genetic closest DNase(+) ST17 strain, as well as between the DNase(+) ST19 1203-05 and the genetic closest DNase(-) ST19 strain.

Overall, a total of 2759 genes were detected after pan-genome analysis within all ST17 strains with assembled genome (n=12), with 1788 genes present in more than 99% of the strains. Only 351 genes were classified as cloud genes (present in less than 15% of strains). A zoom-in analysis focusing the ST17 2211-04 and the closest related 1232-06 strain revealed that the proportion of the shared genome was very high (95%, n=2016/2128 genes), with only 112 genes (5%) differentially present through the 2.1 Mb genome of the two strains. However, 64 of these genes were also present in some of the ST17 DNase(+) strains (and in some of the ST19 strains) leaving us with 48 genes found to be exclusive of the accessory content of DNase(-) ST17 2211-04 (Table S2). Interestingly, these encompass phage-related genes, including genes coding for several hypothetical proteins and for the LexA and the Single-stranded DNA-binding proteins, which are associated with stress response and DNA repair (Erill *et al*., 2007; Boutry *et al*., 2013; Mijakovic *et al*., 2006; Spenkelink *et al*., 2019). In fact, DNase(-) ST17 2211-04 was found to harbor two intact phages of 35Kb (36.7%GC) and 24.6Kb (40.1%GC) integrated in its genome (Table 3). Although the latter is exclusive, not having homology in *S. agalactiae*, it is 96% homolog (99% identity) to *Streptococcus* phage Javan11 (GenBank accession number MK448669.1)

**Table 3.** List of accessory genes differentially present between and within ST17 and ST19 strains.

| Feature | ST17 |  | ST19 |  |
| --- | --- | --- | --- | --- |
|  | DNase(-)<br>2211-04 | DNase(+)<br>(n=14) | DNase(+)<br>1203-05 | DNase(-)<br>(n=17) |
| Phage (24.6Kb) <sup>a</sup> | √ |  |  |  |
| Transposon (TnGBS2.3 homolog) | √ | 776-06 <sup>b</sup> |  |  |
| Cell division protein FtsK |  | √ |  |  |
| Hypothetical protein 1 |  | √ |  |  |
| Hypothetical protein 2 |  | √ |  |  |
| Hypothetical protein 3 |  | √ |  |  |
| Hypothetical protein 4 |  | √ |  |  |
| Serine/threonine-protein kinase PknD |  | √ |  |  |
| Eco47II family restriction endonuclease |  | √ |  |  |
| C-5 cytosine-specific DNA methylase |  | √ |  |  |
| Tyrosine recombinase XerD |  | √ |  |  |
| Laminin-binding protein (Lmb) |  | √ |  | √ |
| Serine-rich repeat protein Srr-1 |  |  | 2 smaller ORFs | √ |
| C5a peptidase |  | √ |  | √ |

<sup>a</sup> Phage (24.6Kb) has no homology in *S. agalactiae* but is 96% homolog (99% identity) to *Streptococcus* phage Javan11 (GenBank accession number MK448669.1)
<sup>b</sup> 96% homology (99.96% identity) with 2211-04

A 34Kb transposon 94% homologous to TnGBS2.3 (Guérillot *et al*., 2013) was also found in DNase(-) ST17 2211-04 strain, carrying several VirB/D4 type IV secretion system (T4SS) and LPXTG protein genes (Figure S3). It is inserted in-frame within the intergenic region (IGR) between the *gbs1087* and *gbs1088* homologs (which code for the fibronectin-binding protein FbsA and the major cardiolipin synthase ClsA, respectively), being flanked by a perfect 13bp direct repetitive sequence (TTACTTTTTAAAT). However, Blastn analysis revealed that this transposon is also present in DNase(+) ST17 776-06 strain. In contrast to 2211-04, the transposon insertion in 776-06 interrupts the transcriptional terminator region of a glycosyltransferase (*gbs0452* homolog), group 2 family protein, involved in the biosynthesis of *S. agalactiae* Group B carbohydrate (Sutcliffe *et al*., 2008). The functional consequences of this insertion are unknown.

Conversely, all ST17 DNase(+) strains were found to uniquely harbor 9 accessory genes (Table 3), the majority of which code for hypothetical proteins (n=4) or are involved in cellular processes and signaling (n=2). Two of these exclusive genes encode the Eco47II family restriction endonuclease and the C-5 cytosine-specific DNA methylase, which are components of prokaryotic DNA restriction-modification mechanisms that protect the organism against invasion by foreign DNA (Roberts *et al*., 2003; Kumar *et al*., 1994). Considering that this accessory gene set is absent in the ST17 DNase(-) 2211-04 as well as in all ST19 strains, it shows up as an important candidate to be involved in the DNases’ activity phenotype, besides the recognized DNase coding genes.

For ST19, the pan-genome analysis detected a total of 3073 genes among all ST19 strains with assembled genome (n=12), of which 1713 were shared by more than 99% of the strains and 878 genes were classified as cloud genes (present in less than 15% of strains). A fined-tune analysis focused on the weak DNase(+) 1203-05 and one of the closest DNase(-) ST19 strains (641-05, 724-06 or 1434-05) revealed that each pair shared at least 93% of the genes (with only 85-130 genes being differentially present). While no accessory content was found to be exclusively present in 1203-05, six genes are uniquely harbored by all ST19 DNase(-) strains (data not shown). However, they are also present in all ST17 strains (including in DNase(-) 2211-04), suggesting that they do not contribute to the DNases’ activity phenotype.

#### 3.6. Virulome characterization

In order to identify putative virulence determinants in *S. agalactiae* that may explain the observed DNase activity exceptions, genomes were queried against a custom database of genes potentially linked to adaptation and pathogenicity. While no specific virulence traits were identified for ST17 DNase(-) 2211-04 when compared with the other ST17 strains, some exclusive particularities were found for ST19 DNase(+) 1203-05 within the ST19 group (Table S3), but it is unlikely that they may contribute for the observed DNase activity phenotype. For instance, both laminin-binding protein (*lmb*) and C5a peptidase (*scpB*) genes, which disrupt complement-mediated innate immunity (Melin, 2011), were found to be absent in the weak DNase(+) strain. However, this is intriguing because the surface C5a peptidase is considered to prevent the recruitment of neutrophils to infection sites and, consequently, preclude the production of NETs (Spellerberg, 2000; Melin, 2011; Pietrocola *et al*., 2018). Additionally, two apparent functional smaller serine-rich repeat protein *srr-1* ORFs were also identified in 1203-05, instead of an intact gene (whose encoded protein is linked to host extracellular matrix) characteristically exhibited by all ST19 strains.

CRISPR-Cas analysis (Figure S4) was not informative in elucidating the disparities of the observed DNase activity phenotypes as it correlates with the core-genome SNV-based phylogeny (Figure S1), with both the DNase(-) 2211-04 and the closest DNase(+) 1232-06 grouping together (cluster 3) in ST17; regarding the weak DNase(+) 1203-05, it confirmed its higher closeness to the related DNase(-) ST19 1434-05, 461-05 and 724-06 strains.

## 4. Discussion

The present work aimed to get insight into the genetic background of DNase production in *S. agalactiae* strains, through WGS-based analyses of clinical strains from carrier and infection, belonging to ST17 and ST19. In 2018, Florindo and co-authors (Florindo *et al*., 2018) found strains displaying a DNase(-) phenotype belonging to a single genetic lineage, CC19, and all CC17 strains displaying a DNase(+) phenotype. As ST17/capsular-type III strains have been associated with LOD and neonatal meningitis (Teatero *et al*., 2016; Martins *et al*., 2017; Almeida *et al*., 2017; Manning *et al*., 2009), these findings seemed to corroborate the hypothesis of a major role of this enzyme in hypervirulence, namely in the evasion processes from the host defense mechanisms (Florindo *et al*., 2018). In the present study, all ST17 strains, except one (DNase(-) ST17 2211-04), were found to be DNase(+), while strains belonging to ST19 were classified as DNase(-), with the exception of strain DNase(+) ST19 1203-05, which revealed some DNase activity. These exceptions in DNase activity from both STs thus constituted an important opportunity to track the genetic basis underlying the DNase production.

While the virulome characterization was not helpful since it did not allow to present genetic characteristics that distinguished each of the exception strains from the ST group they belong to, it was interesting to verify that the analysis of the genetic variability of the seven genes putatively encoding secreted DNases (Derré-Bobillot *et al*., 2013) provided some clues. In fact, we found an exclusive non-synonymous alteration (Glu261Lys) in *gbs0609* for strain DNase(+) ST19 1203-05, although it was not possible to determine the structural effect of this amino acid substitution. On the other hand, a non-synonymous alteration (Thr10Ala) in *gbs0661*, coding for the major *S. agalactiae* nuclease (NucA) (Derré-Bobillot *et al*., 2013), was found in strain DNase(-) ST17 2211-04. Curiously, it involves the substitution of a polar uncharged residue by an amino acid with a hydrophobic side chain within the hydrophobic domain of the predicted signal peptide, which is required for export/secretion of the mature protein (Paetzel *et al*., 2002). Although an hydrophobicity increase was showed to favor protein processing and translocation (Owji *et al*., 2018; Paetzel *et al*., 2002), we observed an opposite phenotypic scenario, so we speculate whether the observed mutation affecting NucA signal peptide may have impaired the protein secretion, leading to DNase(-) phenotype. In another perspective, we did not found any difference in the predicted NucA mature protein among all ST17 and all ST19 strains, regardless of the DNase activity phenotype. When compared to the well-studied NEM316 NucA, they only differ at one position (Pro99Leu) (as the additional nucleotide variation seen is synonymous), containing a conserved H-N-N motif (H^148^, N^167 or 170^ and N^179^) and other key amino acid residues (like Glu^193^) that are crucial for enzyme activity and metal binding (Moon *et al*., 2014). Therefore, the allelic difference between all these proteins and the NEM316 NucA mature protein seems not justify any differences in the DNase activity phenotype, since some ST17 strains presented a strong DNase activity, similar to NEM316. Accordingly, it is very likely that, besides NucA and other predicted DNases, other proteins, namely those involved in their export/secretion, may account for the DNase phenotypic differences observed between ST17 and ST19 strains.

The analysis of the core-genome identified some specificities (SNVs or indels) differentiating the DNase(-) ST17 2211-04 and the DNase(+) ST19 1203-05, from all DNase(+) ST17 strains and all DNase(-) ST19 strains, respectively. We thus looked for a rationale that could justify why some of the altered genes could potentially be implicated in DNase release or activity. For instance, ATP-dependent Clp protease ATP-binding subunit ClpC and ClpX, the molecular chaperone DnaK and the class I SAM-dependent methyltransferase were mutated in the DNase(-) ST17 2211-04 and are involved in metabolism, stress response and pathogenesis (Frees *et al*., 2004; Nair *et al*., 2003; Aguilar-Rodríguez *et al*., 2016; Chiappori *et al*., 2015; Tomoyasu *et al*., 2012; Grove *et al*., 2011; Kozbial and Mushegian, 2005; Struck *et al*., 2012). Genes related to bacterial stress response and DNA repair were mutated in DNase(+) ST19 1203-05, namely genes encoding for the GTPase Obg and the excinuclease ABC subunit B. GTPase Obg has been associated with a variety of cellular functions, including regulation of the cell stress response and persistence in response to nutrient starvation, while the excinuclease ABC subunit B is pivotal for stress responses, and recognition and processing of DNA lesions (Bonventre *et al*., 2016; Grossman and Yeung, 1990). For DNase(+) ST19 1203-05 strain, the gain of a premature stop codon in the TetR/AcrR family transcriptional regulator gene could play an important role in metabolic regulation. This family of transcriptional regulators is widely associated with antibiotic resistance and with the regulation of genes encoding small-molecule exporters (Ramos *et al*., 2005; Cuthbertson and Nodwell, 2013; Deng *et al*., 2013). Moreover, the ST19 DNase(+) strain presented a core-SNV within a non-coding sequence of a threonine/serine exporter family protein, which hypothetically may influence the expression profile. Interestingly, serine/threonine signaling cascades were linked with virulence, namely in *S. agalactiae* (Rantanen *et al*., 2007). However, the impact of these mutations in DNase enzymatic performance remains undetermined. In fact, enzymes act under of multiple levels of regulation and, in order to clarify the overall picture of these regulation processes, different levels of “omics” data such as transcriptomics, proteomics, metabolomics and epigenomics should be integrated (Shimizu, 2013; Willenborg and Goethe, 2016).

Concerning the pan-genome, the analysis was focused on genes exclusive of the ‘exception strains’ for DNase activity (DNase(-) ST17 2211-04 and DNase(+) ST19 1203-05). A total of 48 phage-related genes were found to be exclusive of DNase(-) ST17 2211-04. Although it was not possible to determine the exact integration coordinates in the genome, this strain was found to harbor a unique intact phage with no homology in available *S. agalactiae* genome sequences. In a pure speculative basis, this could somehow contribute to the DNase phenotype of this strain. In fact, phages were found to regulate bacterial populations by altering bacterial gene expression, as well as to contribute for the evolution of bacterial hosts (through gene transfer) and to bacterial pathogenesis (Mee-Marquet *et al*., 2018; Javan *et al*., 2019). The pan-genomic analysis also allowed the identification of a transposon homologous to TnGBS2.3 in two ST17 strains (DNase(-) 2211-04 and DNase(+) 776-06), yet with different sites of insertion. TnGBS transposons are common among *S. agalactiae* isolates and carry several VirB/D4 T4SS and LPXTG protein genes (Guérillot *et al*., 2013). While VirB/D4 T4SS mediates the conjugative transfer of DNA, enhances bacterial pathogenicity and was proposed to mediate largescale infectious outbreaks in humans (Guérillot *et al*., 2013; Zhang *et al*., 2012), LXPTG have been shown to be involved in host colonization, biofilm formation and immune modulation (Brady *et al*., 2010; Maddocks *et al*., 2011). The site of insertion of the transposon in DNase(-) ST17 2211-04 strain is proximal to genes belonging to a specialized system for excreting extracellular proteins across bacterial cell membranes, the type VII protein secretion system, which has been associated with virulence in *Staphylococcus aureus* (Warne *et al*., 2016). Although the transference of new genes to *S. agalactiae* genome is thought to modify the expression of neighboring genes at the integration site (Fléchard and Gilot, 2014), appropriate transcriptomic analysis is required to evaluate the impact of this transposon insertion.

The identification of nine accessory genes among all ST17 DNase(+) strains could also be related with the phenotypes observed for DNase production. Among these genes there are two encoding for proteins involved in mechanisms that protect the organism against invasion by foreign DNA (Roberts *et al*., 2003), the Eco47II family restriction endonuclease and the C-5 cytosine-specific DNA methylase. The observation that none of these loci was found in any DNase(-) strain may suggest a contribution of these proteins to the lack of DNase activity. Nevertheless, this hypothesis also warrants further investigation through functional assays.

In conclusion, we globally characterized the genome of *S. agalactiae* ST17 and ST19 strains, isolated from invasive and carriage clinical presentations, DNase(+) and DNase(-), in an attempt to shed some light on the genetic background underlying *S. agalactiae* DNase activity. Some loci and SNVs / indels potentially contributing to DNase activity were identified. However, these data require further research to test their robustness, starting by evaluating *S. agalactiae* strains of different ST and diverse DNase production phenotypes, in order of reaching a better understanding of the pathways whereby DNases contribute to *S. agalactiae* pathogenesis.

## Acknowledgments

The authors wish to thank Barbara Spellerberg for the kind gift of *S. agalactiae* strains from collection of the Institute of Medical Microbiology and Hygiene (IMMH), Ulm University, Germany, and to Ana Tenreiro of the Faculdade de Ciências of University of Lisbon, for the use of the fluorimeter. This work was supported by the GenomePT project (POCI-01-0145-FEDER-022184) from Fundação para a Ciência e a Tecnologia (FCT).

