## Supplementary figures and images for "Genomic insights on DNase production in *Streptococcus agalactiae* ST17 and ST19 strains"

### Figure S1

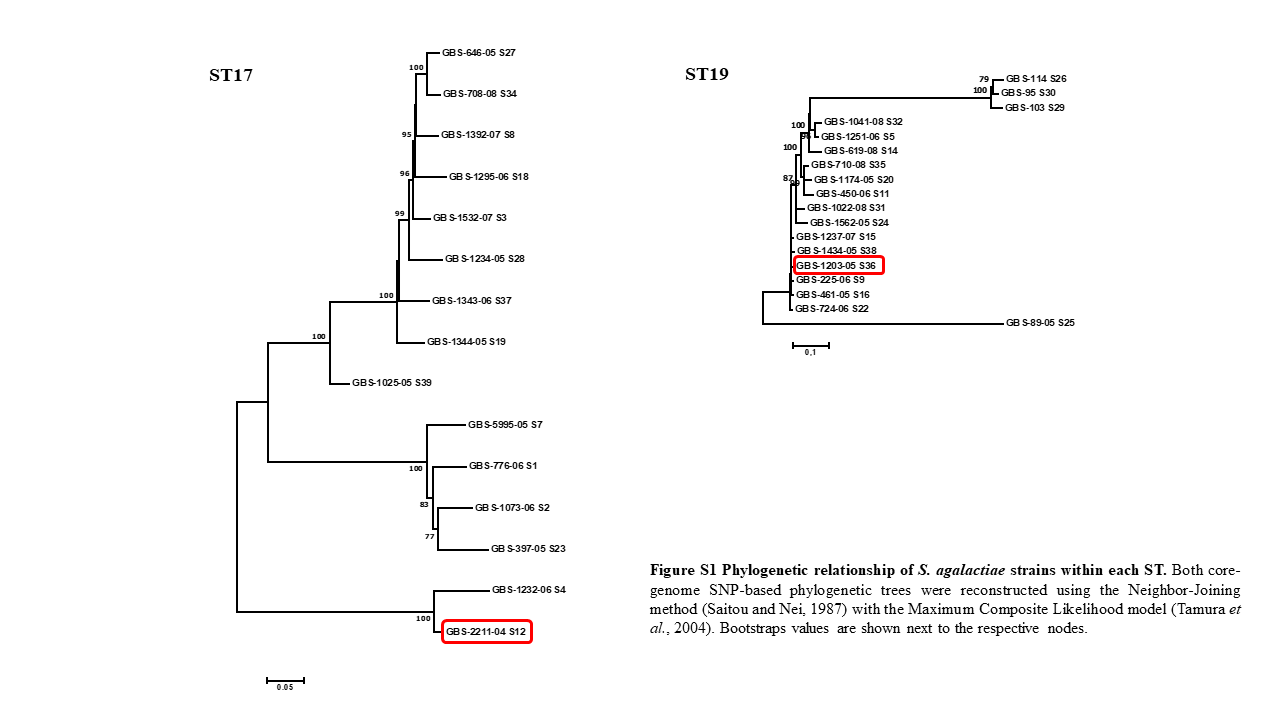

### Figure S2

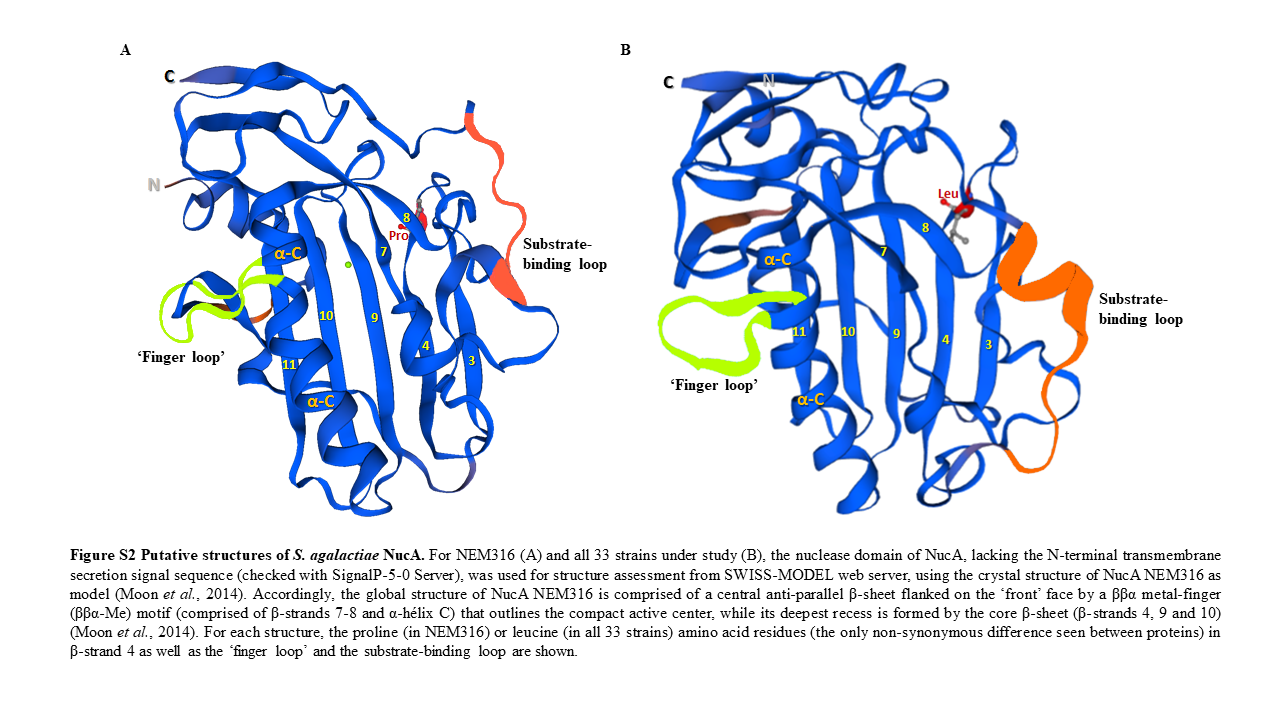

### Figure S3

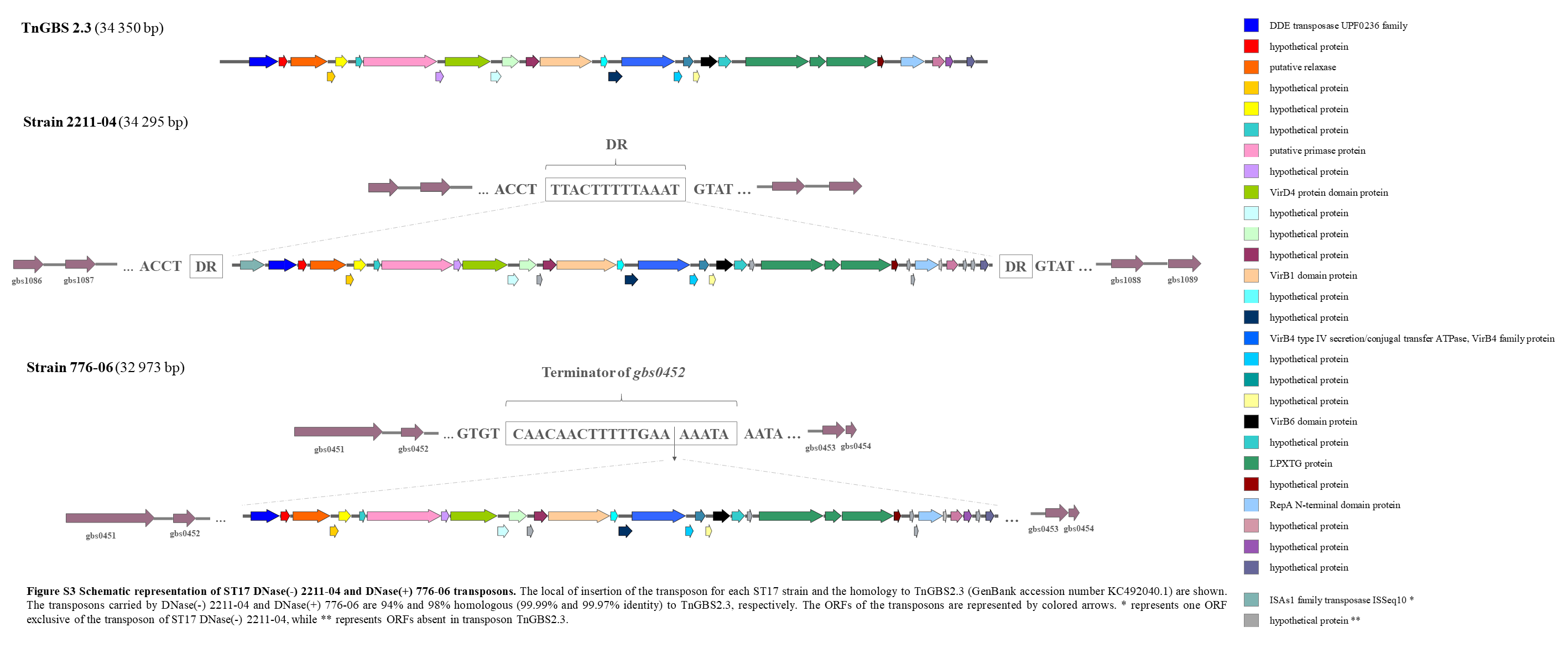

### Figure S4

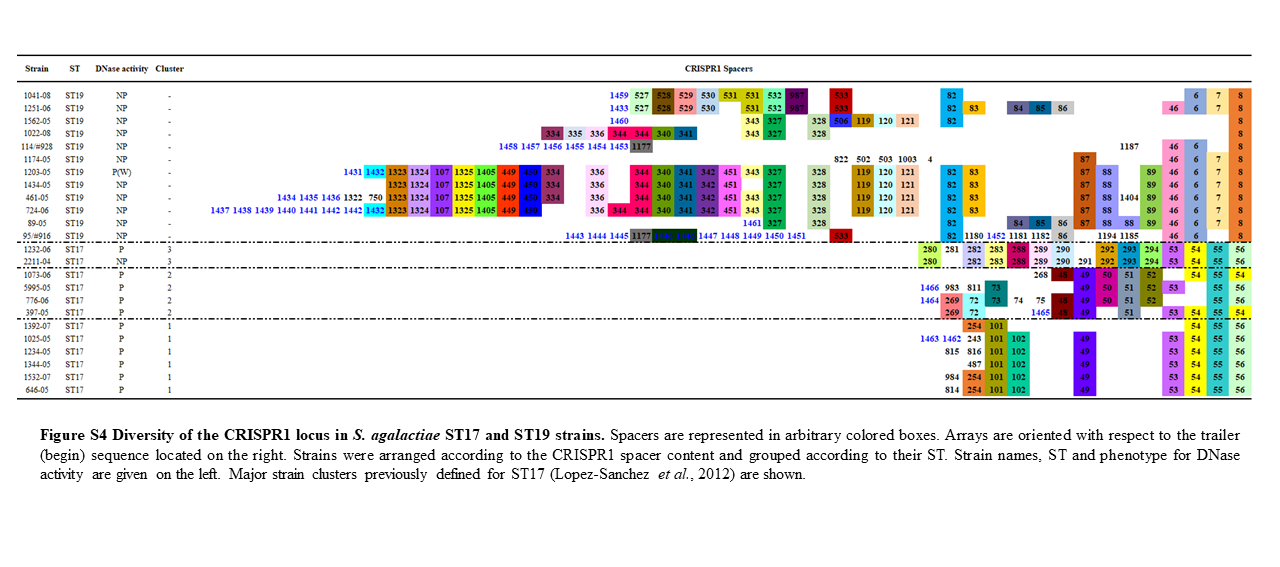
