## Supplementary material for "Genomic insights on DNase production in *Streptococcus agalactiae* ST17 and ST19 strains": Table S1

**Table S1** *S. agalactiae* human strains data information.

| Isolate ID | Isolation Country | Year | Isolation Source | Host Gender | Host Age | Disease | ENA Accession Sample | ENA ID |
|---|---|---|---|---|---|---|---|---|
| 1022-08 | Portugal | 2008 | vaginal/rectal | female | 33 | carrier | ERS5323489 | PT_GBS0001 |
| 1025-05 | Portugal | 2005 | vaginal/rectal | female | 29 | carrier | ERS5323490 | PT_GBS0002 |
| 103/#923 | Germany | 2001-2003 | blood/CSF | unknown | < 3 months | invasive | ERS5323491 | PT_GBS0003 |
| 1041-08 | Portugal | 2008 | vaginal/rectal | female | 38 | carrier | ERS5323492 | PT_GBS0004 |
| 1073-06 | Portugal | 2006 | vaginal/rectal | female | 17 | carrier | ERS5323493 | PT_GBS0005 |
| 114/#928 | Germany | 2001-2003 | blood/CSF | unknown | < 3 months | invasive | ERS5323494 | PT_GBS0006 |
| 1174-05 | Portugal | 2005 | vaginal/rectal | female | 22 | carrier | ERS5323495 | PT_GBS0007 |
| 1203-05 | Portugal | 2005 | vaginal/rectal | female | 21 | carrier | ERS5323496 | PT_GBS0008 |
| 1232-06 | Portugal | 2006 | vaginal/rectal | female | 22 | carrier | ERS5323497 | PT_GBS0009 |
| 1234-05 | Portugal | 2005 | vaginal/rectal | female | 34 | carrier | ERS5323498 | PT_GBS0010 |
| 1237-07 | Portugal | 2007 | vaginal/rectal | female | 29 | carrier | ERS5323499 | PT_GBS0011 |
| 1251-06 | Portugal | 2006 | vaginal/rectal | female | 71 | carrier | ERS5323500 | PT_GBS0012 |
| 1295-06 | Portugal | 2006 | vaginal/rectal | female | 35 | carrier | ERS5323501 | PT_GBS0013 |
| 1343-06 | Portugal | 2006 | vaginal/rectal | female | 35 | carrier | ERS5323502 | PT_GBS0014 |
| 1344-05 | Portugal | 2005 | vaginal/rectal | female | 20 | carrier | ERS5323503 | PT_GBS0015 |
| 1392-07 | Portugal | 2007 | vaginal/rectal | female | 25 | carrier | ERS5323504 | PT_GBS0016 |
| 1434-05 | Portugal | 2005 | vaginal/rectal | female | 31 | carrier | ERS5323505 | PT_GBS0017 |
| 1532-07 | Portugal | 2007 | vaginal/rectal | female | 30 | carrier | ERS5323506 | PT_GBS0018 |
| 1562-05 | Portugal | 2005 | vaginal/rectal | female | 20 | carrier | ERS5323507 | PT_GBS0019 |
| 2211-04 | Angola | 2004 | CSF | male | < 12 years | meningitis | ERS5323508 | PT_GBS0020 |
| 225-06 | Portugal | 2006 | vaginal/rectal | female | 25 | carrier | ERS5323509 | PT_GBS0021 |
| 397-05 | Portugal | 2005 | vaginal/rectal | female | 29 | carrier | ERS5323510 | PT_GBS0022 |
| 450-06 | Portugal | 2006 | vaginal/rectal | female | 24 | carrier | ERS5323511 | PT_GBS0023 |
| 461-05 | Portugal | 2005 | vaginal/rectal | female | 20 | carrier | ERS5323512 | PT_GBS0024 |
| 5995-05 | Angola | 2005 | CSF | male | < 12 years | meningitis | ERS5323513 | PT_GBS0025 |
| 619-08 | Portugal | 2008 | vaginal/rectal | female | 49 | carrier | ERS5323514 | PT_GBS0026 |
| 646-05 | Portugal | 2005 | vaginal/rectal | female | 33 | carrier | ERS5323515 | PT_GBS0027 |
| 708-08 | Portugal | 2008 | vaginal/rectal | female | 23 | carrier | ERS5323516 | PT_GBS0028 |
| 710-08 | Portugal | 2008 | vaginal/rectal | female | 41 | carrier | ERS5323517 | PT_GBS0029 |
| 724-06 | Portugal | 2006 | vaginal/rectal | female | 25 | carrier | ERS5323518 | PT_GBS0030 |
| 776-06 | Portugal | 2006 | vaginal/rectal | female | 59 | carrier | ERS5323519 | PT_GBS0031 |
| 89-05 | Portugal | 2005 | vaginal/rectal | female | 26 | carrier | ERS5323520 | PT_GBS0032 |
| 95/#916 | Germany | 2001-2003 | blood/CSF | unknown | < 3 months | invasive | ERS5323521 | PT_GBS0033 |

ID, Identification; CSF, Cerebrospinal Fluid
