## Supplementary material for "Genomic insights on DNase production in *Streptococcus agalactiae* ST17 and ST19 strains": Table S2

**Table S2** List of accessory genes exclusive of DNase(-) ST-17 2211-04 strain.

| Product | |
|---|---|
| Phage integrase | Hypothetical protein |
| Hypothetical protein | Hypothetical protein |
| LexA repressor | Phage protein |
| Transcriptional regulator | Phage terminase, small subunit |
| Phage antirepressor protein | Phage terminase, large subunit |
| Hypothetical protein | Phage minor capsid protein |
| Hypothetical protein | Phage minor capsid protein |
| Hypothetical protein | Hypothetical protein |
| Hypothetical protein | Hypothetical protein |
| Hypothetical protein | Hypothetical protein |
| Hypothetical protein | Hypothetical protein |
| Hypothetical protein | Hypothetical protein |
| Phage recombination protein Bet | Hypothetical protein |
| Phage protein | Phage protein |
| Hypothetical protein | Putative minor capsid protein - phage associated |
| Hypothetical protein | Phage minor capsid protein |
| Single-stranded DNA-binding protein | Phage minor capsid protein |
| Phage protein | Phage major tail shaft protein |
| Hypothetical protein | hypothetical protein |
| Phage protein | Phage protein |
| Phage protein | Hypothetical protein |
| Hypothetical protein | Hypothetical protein |
| Phage protein | Paratox |
| Hypothetical protein | Phage lysin, N-acetylmuramoyl-L-alanine amidase |
