## Supplementary material for "Genomic insights on DNase production in *Streptococcus agalactiae* ST17 and ST19 strains": Table S3

**Table S3** List of virulence factors differently present in DNase(+) ST19 1203-05.

| | | | ST17 clinical strains | | ST19 clinical strains | |
|---|---|---|---|---|---|---|
| **Virulence Factors** | **NEM316 old locus tag** [a] | **2603V/R locus tag** [b] | **DNase(-) 2211-04** | **DNase(+) (n=14)** | **DNase(+) 1203-05** | **DNase(-) (n=17)** |
| Laminin-binding protein (Lmb) | gbs1307 | SAG1234 | √ | √ | --- | √ |
| Serine-rich repeat protein Srr-1 | --- | SAG1462 | --- | --- | 2 smaller ORFs | √ |
| C5a peptidase | gbs1308 | SAG1236 | √ | √ | --- | √ |

[a] ORF designations according to NEM316 genome (GenBank accession number NC004368.1)
[b] ORF designations according to 2603V/R genome (GenBank accession number NC004116.1)
